# Testing for genetic assimilation with phylogenetic comparative analysis: Conceptual, methodological, and statistical considerations

**DOI:** 10.1101/2021.12.28.473512

**Authors:** Alex R. Gunderson, Liam J. Revell

## Abstract

Genetic assimilation is a process that leads to reduced phenotypic plasticity during adaptation to novel conditions, a potentially important phenomenon under global environmental change. Null expectations when testing for genetic assimilation, however, are not always clear. For instance, the statistical artifact of regression to the mean could bias us towards detecting genetic assimilation when it has not occurred. Likewise, the specific mechanism underlying plasticity expression may affect null expectations under neutral evolution. We used macroevolutionary numerical simulations to examine both of these important issues and their interaction, varying whether or not plasticity evolves, the evolutionary mechanism, trait measurement error, and experimental design. We also modified an existing reaction norm correction method to account for phylogenetic non-independence. We found: 1) regression to the mean is pervasive and can generate spurious support for genetic assimilation; 2) experimental design and post-hoc correction can minimize this spurious effect; and 3) neutral evolution can produce patterns consistent with genetic assimilation without constraint or selection, depending on the mechanism of plasticity expression. Additionally, we re-analyzed published macroevolutionary data supporting genetic assimilation, and found that support was lost after proper correction. Considerable caution is thus required whenever investigating genetic assimilation and reaction norm evolution at macroevolutionary scales.

## Introduction

Phenotypic plasticity is an important phenomenon in evolutionary biology with far-reaching implications (Ghalambor et al. 2007; Murren et al. 2015). For example, plasticity in a given trait can facilitate evolutionary radiations (Pfennig et al. 2010), contribute to local adaptation (Price et al. 2003), and dampen selection on underlying genetic variation (Price et al. 2003; Crispo 2008). Interest in the expression and evolution of phenotypic plasticity has increased in recent years due to the intense selection pressure that human activity has imposed on the natural world (Chevin et al. 2010; Kingsolver and Buckley 2017). Taxa with greater plasticity in traits that are important for tolerating human-induced changes to the environment are predicted to fare better under the novel conditions created by global warming, urbanization, and species introductions (Somero 2010; Gunderson and Stillman 2015; Seebacher et al. 2015; Gunderson et al. 2017). For instance, plasticity in thermal physiology is predicted to decrease the negative impact of global warming on ectotherms (Gunderson et al. 2017; Riddell et al. 2018; Rohr et al. 2018; Morley et al. 2019). Greater knowledge of the forces that shape and constrain phenotypic plasticity is crucial both for our fundamental understanding of ecology and evolution and our ability to predict and mitigate the consequences of global change.

Adaptation to novel conditions is sometimes associated with a reduction or loss of plasticity in traits under selection, a process that is known as genetic assimilation (Waddington 1953; Schlichting and Pigliucci 1998). Some have argued that genetic assimilation is a key mechanism of adaptive evolutionary change (West-Eberhard 2003; Crispo 2007). For example, the canalization of formerly induced phenotypes can provide a means of adaptive phenotypic divergence during local adaptation and adaptive radiation (Ehrenreich and Pfennig 2016; Martin et al. 2016; Gunter et al. 2017; Schneider and Meyer 2017). In addition, genetic assimilation may be an outcome of population responses to anthropogenic global change. Lande (2009) found that the evolution of increased phenotypic plasticity followed by genetic assimilation should result from selection due to dramatic changes in the environment.

The intersection of adaptation, genetic assimilation, and global change is highlighted by the Baseline Tolerance/Tolerance Plasticity Trade-off Hypothesis (hereafter referred to as the Trade-off Hypothesis). Emerging from the field of evolutionary physiology, the hypothesis states that, as organisms evolve greater baseline tolerance to extreme temperatures (heat or cold), plasticity in their thermal tolerance should actually decrease (van Heerwaarden and Kellermann 2020). In other words, high levels of constitutive thermal tolerance evolve by genetic assimilation, resulting in thermal tolerance phenotypes that are less sensitive to environmental temperature variation (Sikkink et al. 2014). Support for this hypothesis comes from a variety of organisms, including marine and aquatic vertebrates and invertebrates at both the intraspecific and interspecific levels (e.g., Stillman 2003; Esperk et al. 2016; Comte and Olden 2017; Armstrong et al. 2019). One of several implications of the Trade-off Hypothesis is that the genetic mechanisms associated with the expression of physiological plasticity serve as the raw material for thermal adaptation. Therefore, as organisms evolve greater baseline tolerance in response to global change, they may lose both the ability to evolve even greater tolerance and their capacity to track shorter-term environmental changes via a plastic response (van Heerwaarden and Kellermann 2020).

Hypotheses that make predictions about trait change over time among individuals and/or taxa, such as the Trade-off Hypothesis, are prone to spurious support due to the well-known statistical phenomenon called *regression to the mean*. A consequence of this phenomenon is that when a particular trait is measured repeatedly, the experimental units (e.g., species, populations, or individuals) that start with particularly high or low values relative to the population mean are also likely to record the greatest change in value over time as subsequent measures regress towards the mean (Barnett et al. 2005). This is not a real biological effect: rather, it is an artifact created by sampling and measurement error.

Regression to the mean is a common problem in repeated-measures analyses, and statistical methods have been developed to overcome the spurious results that it can create. For example, data resampling techniques have been applied to generate null distributions of trait change (Jackson and Somers 1991), and methods to statistically remove regression to the mean effects prior to statistical analysis have also been proposed (Kelly and Price 2005). These approaches are most often employed at the intraspecific level (Ghalambor et al. 2015; Deery et al. 2021), but accounting for regression to the mean is also necessary, though more often neglected, in interspecific macroevolutionary analyses (Baker et al. 2015).

Motivated by current debate surrounding the Trade-off Hypothesis and increasing interest in the evolution of phenotypic plasticity and genetic assimilation under global change (Gunderson and Stillman 2015; Kelly 2019; Sasaki and Dam 2019; van Heerwaarden and Kellermann 2020; Barley et al. 2021; Preston et al. 2021; Sasaki and Dam 2021), we used simulations to explore the more general consequences of regression to the mean for tests of genetic assimilation at the macroevolutionary level. We conducted simulations that assumed mechanistically different models for plasticity evolution, as well as different experimental sampling designs for measuring species phenotypic traits. In addition, we conducted simulations with and without the assumption that phenotypes are measured with error. We found that measurement error alone can be sufficient to recover a macroevolutionary pattern that is consistent with genetic assimilation due entirely to the phenomenon of regression to the mean. This problem can be avoided, however, by using an appropriate experimental design. In addition, we found that macroevolutionary patterns consistent with genetic assimilation are the null expectation under random Brownian motion evolution for some models by which plasticity evolves. Finally, we applied a statistical method designed to remove the regression to the mean effect from compromised data (Kelly and Price 2005), that we modified to take non-independence due to phylogeny into account. We found that our approach can reduce the effect of regression to the mean in some, but not all, circumstances. We applied this modified approach to published phylogenetic comparative data that was thought to be consistent with the Trade-off Hypothesis, and showed that the relationship is not significant when properly corrected for regression to the mean.

## Methods

In our simulations, we assumed that there were two different environments that induce different phenotypes. The difference between trait values in each environment represented the magnitude of phenotypic plasticity and, in a two-environment system, is equivalent to the slope of the reaction norm. In any given simulation, we first generated a random phylogenetic tree with 50 terminal nodes using the *pbtree* function in the phytools R package (Revell 2012; R Development Core Team 2021). Depending on the simulation, we either modeled the evolution of plasticity across the phylogeny using the *fastBM* function or assumed no phenotypic evolution at all (see below for details). In addition, we assumed a certain degree of sampling error in the measurement of phenotypes in each environment. We did so by drawing phenotypes for individuals randomly from a Gaussian distribution with a mean value at the true phenotypic population mean. In practice, this type of error can emerge both from the random sampling of individuals with a population that varies, and by methodological or instrument imprecision. Since both types of error can contribute to the phenomenon of regression to the mean, we effectively combined them in our simulation. For each simulation run, we calculated the phylogenetic correlation coefficient for the association between the baseline phenotype and the reaction norm slope using the *phyl*.*vcv* function of phytools. Starting conditions assumed a baseline phenotypic trait value of 0 and a phenotypic value in the novel environment of 1. In simulations where evolution occurred, the Brownian motion rate parameter (*σ*^2^) was set at 1, 0.1, or 0.01 for respective traits. Phenotypic measurement error was set to have variances (*s*^2^) of 1, 0.1, or 0.01.

We predicted that experimental design would influence whether or not regression to the mean biased our results. As such, we modeled three different experimental designs for measuring the phenotypes of each taxon in each environment (Figure 1). In Design 1, we assumed that each individual has their phenotype measured twice in total: once after acclimation to each environment. Therefore, in this case the same individuals are used to estimate both the baseline phenotype and the reaction norm. In Design 2, we assumed that different individuals are acclimated to each environment prior to phenotypic measurement. In this case, individuals exposed to the baseline environment are used to estimate the baseline phenotype, and the difference in phenotype between the two sets of individuals is used to estimate the reaction norm. Finally, in Design 3, we assumed that one set of individuals is acclimated only to the baseline environment prior to phenotypic trait measurement, and that a second set of individuals have their phenotypes measured twice: once after acclimation to each of the two environments. The former individuals are used only to estimate baseline phenotypes, and the latter individuals are used only to estimate the slope of the reaction norm.

**Figure 1.**
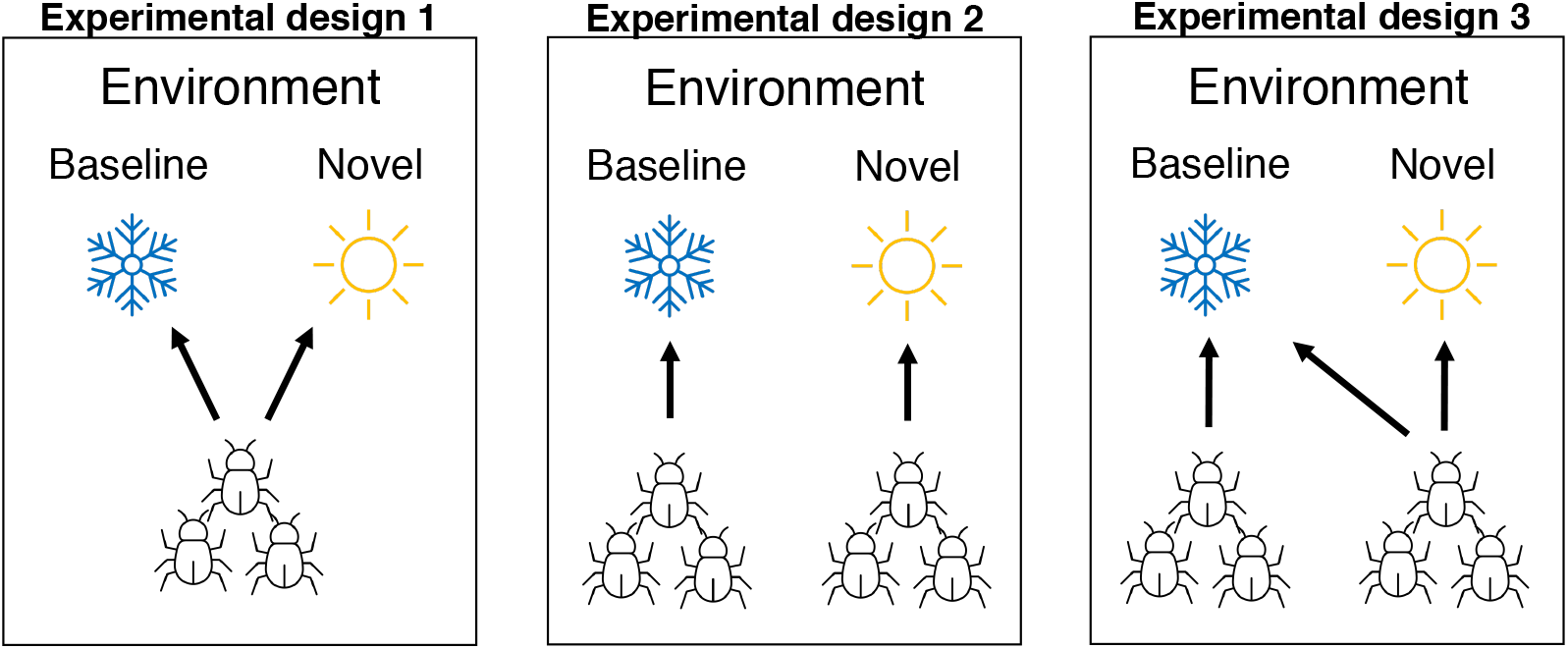
Three experimental designs that we simulated for estimating baseline phenotype and phenotypic plasticity of species. In Experimental Design 1, baseline phenotype and plasticity are estimated from a single group of individuals that have their phenotype measured twice: once after exposure to each environment. In Experimental Design 2, there are two groups of individuals that have their phenotype measured only once, after exposure to either the baseline or novel environment. In Experimental Design 3, there are also two groups of individuals. One group has their phenotype measured only after exposure to the baseline condition, and are used only to estimate the baseline phenotype. The other group of individuals have their phenotypes measured twice (as in Experimental Design 1), but they are only used to estimate phenotypic plasticity. Thermal environments are shown simply as an example.

We also tested whether regression to the mean could emerge as a significant problem when phenotypic plasticity evolves, and did so assuming a neutral evolutionary process (in this case, Brownian motion). We used two different models for the evolution of phenotypic plasticity, that we refer to as Linked Phenotypes and Unlinked Phenotypes. In the Linked Phenotypes model, we assumed that the slope and intercept of the reaction norms evolve directly, with the phenotype that is expressed in each environment evolving indirectly as a byproduct. This is one of the most common ways in which the evolution of phenotypic plasticity is conceptualized and studied theoretically (Lande 2009; Chevin et al. 2010; Chevin et al. 2013). We refer to this as the Linked Phenotypes model because the phenotypes that are expressed in each environment are each determined by the slope and the intercept, linking them under a neutral evolutionary process. In the Unlinked Phenotypes model, the phenotype that is expressed in a given environment evolves directly and independently of the phenotype expressed in the other environment. Therefore, the phenotypes that are expressed in each environment lack genetic covariance (Via and Lande 1985), and the slope and intercept of the reaction norm emerge merely as a byproduct of the trait values expressed in each environment. This model is based on the concept that parts of a reaction norm can evolve independently of one another as outlined in Ghalambor et al. (2007). Using each model, we conducted simulations in which the evolving parameters diverge via Brownian motion, and then estimated the phylogenetic correlation coefficient for an association between baseline phenotype and the reaction norm slope, as described above.

When possible, we applied a statistical correction developed by Kelly and Price (2005) to remove the regression to the mean effect from reaction norm values. The method was not originally designed for use with phylogenetic comparative data, but we modified it to that purpose here. We use the correction only with data simulated under Experimental Designs 1 and 2, as the method cannot be applied to Experimental Design 3. The approach is based on the premise that if the plastic response is constrained by the baseline phenotype, then phenotypic variation should be reduced under novel conditions and that phenotypes in the baseline and novel conditions should be correlated within experimental units (Kelly and Price 2005). The approach is implemented as follows. First, we estimate the phylogenetic variance in phenotypes in each environment (*s*_*i*_) using the *phyl*.*vcv* function, then we conduct a Pittman’s test of homogeneity of variance. If there is no difference in variance between the phenotypes, then corrected reaction norms 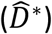 are calculated using the following function:

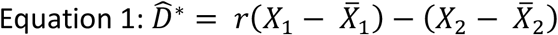

where *r* is the phylogenetic correlation coefficient for the relationship between phenotypes in the baseline and novel environments, *X*_*i*_ is the vector of phenotypes in a given environment, and 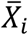 is the phylogenetic phenotypic mean in a given environment. If the phenotypic variance differs between environments, then an adjusted correlation coefficient 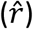 is used in Equation 1, calculated as follows:

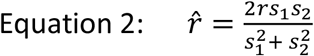

Where 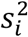 is the estimated phylogenetic variance in the environment *i*. 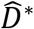 is then used as the reaction norm value for subsequent analyses as described above.

Using our modified Kelly and Price (2005) approach, we reanalyzed previously published data on heat tolerance plasticity across Nudibranch species (Armstrong et al. 2019). The authors of that study used what we refer to as Experimental Design 2 (Figure 1), and originally found support for the Trade-off Hypothesis, as Nudibranch species with greater baseline heat tolerance had reduced heat tolerance plasticity. However, they did not adjust values for potential regression to the mean as we do here.

## Results

### Results without evolution but with measurement error

We first present results of simulations in which phenotypic plasticity is present but does not diverge among taxa. In this case, the only source of variation among taxa is measurement error. If regression to the mean is not an issue, we would expect the correlation coefficients generated in our simulations to be centered on 0.

Instead, we found that measurement error generated a pattern consistent with genetic assimilation (that is, a negative correlation between baseline phenotype and reaction norm slope) due to regression to the mean, but only for Experimental Designs 1 and 2 (Figure 2). By contrast, regression to the mean does not bias results using Experimental Design 3 (Figure 2). This finding does not dependent on the level of plasticity organisms exhibit. For example, even if there is no plasticity in the trait, a negative relationship between baseline phenotype and reaction norm slope will emerge with Experimental Designs 1 and 2 if the phenotypes in each environment are measured with error (see Supplemental Figure S1; see also Deery et al. 2001 for an individual-level example). The bias induced by Experimental Designs 1 and 2 is largely removed, however, when the simulated data are adjusted using our phylogenetic modification of the Kelly and Price (2005) correction (Figure 2 “corrected estimates”).

**Figure 2.**
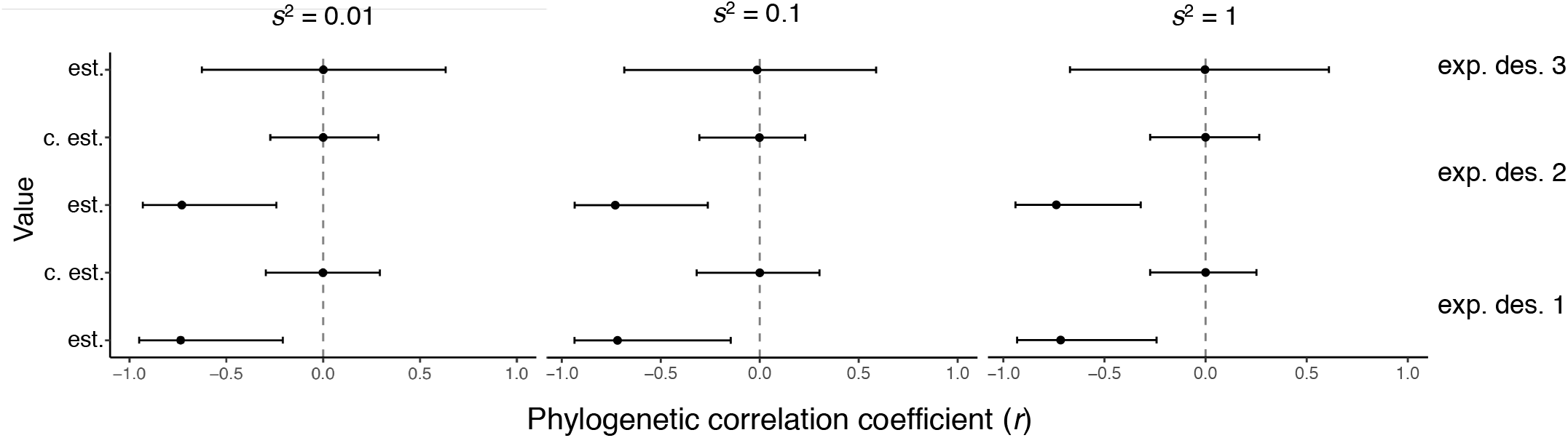
Summary of phylogenetic correlation coefficients between baseline phenotype and reaction norm slope from simulations in which plasticity is present but does not diverge among taxa, and phenotypes in each environment are measured with error. Points indicate the median and black lines the central 95% of correlation coefficients from 1000 simulations. The error variance of simulations is shown above each column. Alternating shading differentiates different simulated experimental designs, which are written on the righthand side of the figure. “est.” denotes estimated correlation coefficients given measurement error, while “c. est” denotes those same correlation coefficients adjusted to remove the effect of regression to the mean following our modified Kelly and Price (2005) correction. Adjusted results are not given for Experimental Design 3 because the adjustment cannot be applied to data collected with that design.

### Results with neutral evolution of phenotypic plasticity

A pattern consistent with genetic assimilation is unlikely to evolve under neutral processes if phenotypic plasticity evolves via the Linked Phenotype mechanism (i.e., if reaction norm slope and intercept evolve directly) (Figure 3 “true” values). However, this assumes that there is no measurement error when estimating phenotypes. When measurement error is considered, regression to the mean can produce spurious support for genetic assimilation with Experimental Designs 1 and 2, though this effect depends on the magnitude of measurement error relative to the rate of evolution (Figure 3 “estimates” values). The greater the measurement error, the more likely that regression to the mean can become a problem.

**Figure 3.**
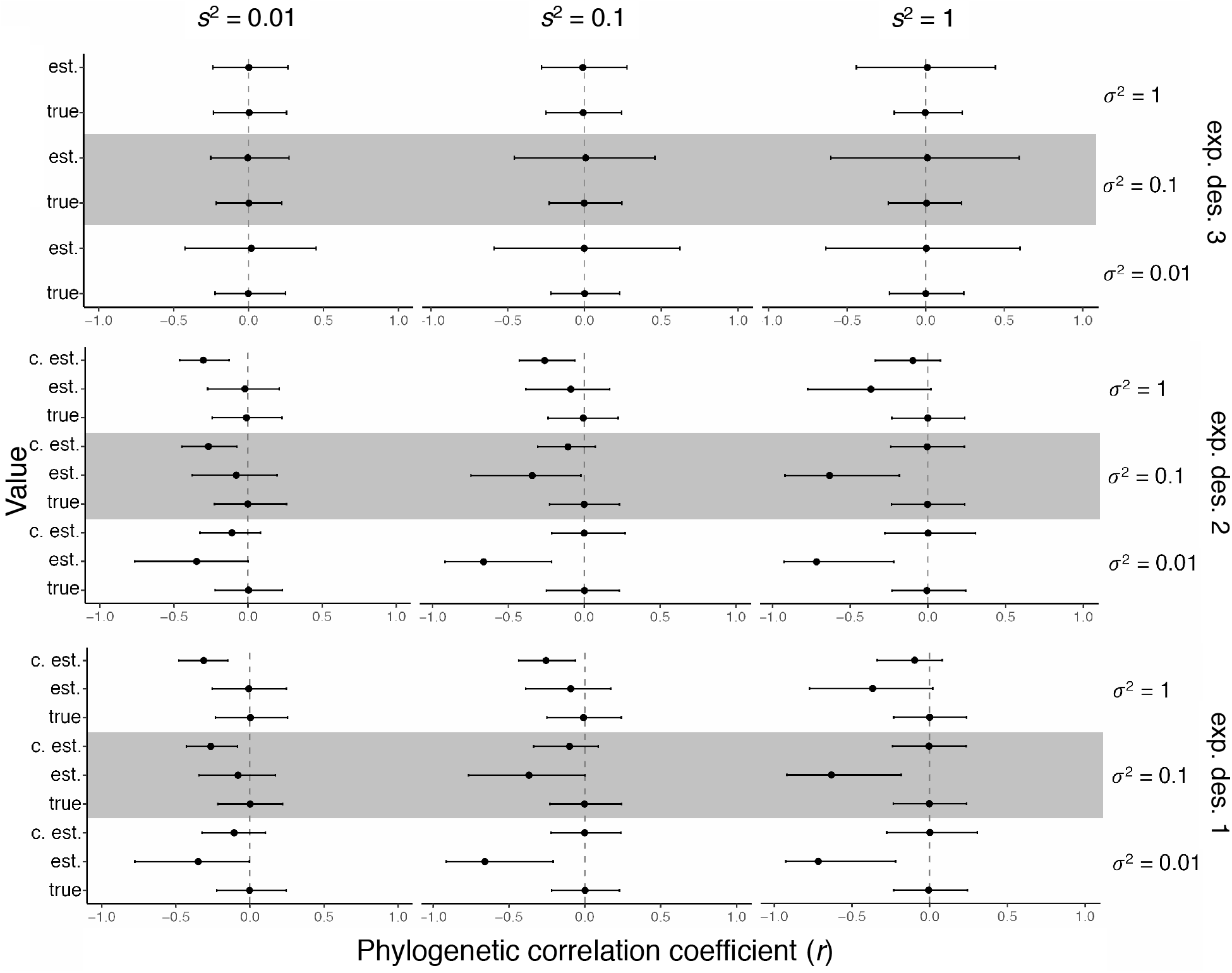
Summary of phylogenetic correlation coefficients between baseline phenotype and reaction norm slope from simulations in which plasticity evolved by Brownian motion via the Linked Phenotypes mechanism. Each panel shows the result of 1000 simulations. Each row shows the results of simulations with a different experimental design (Figure 1): *Bottom*, Design 1. *Middle*: Design 2. *Top*: Design 3. Alternating shaded areas differentiate simulations with different Brownian motion rate parameters (*σ*^2^). Each column contains simulations with a different error variance for phenotypic estimation (*s*^2^). “true” denotes the true phylogenetic correlation values from the simulations. “est.” denotes estimates of the true correlation coefficients given measurement error. “c. est” denotes estimated correlation coefficients corrected to remove the effect of regression to the mean following our modified Kelly and Price (2005) adjustment. Adjusted results are not given for Experimental Design 3 because the method cannot be applied to data collected with that design.

Adjusting data generated by Experimental Designs 1 and 2 using our modified Kelly and Price correction removes the regression to the mean effect (Figure 3 “corrected estimates”). These results, however, also highlight the importance of judicious application of the Kelly and Price correction. This is because if there is a weakly negative to positive correlation between baseline phenotype and plasticity, the Kelly and Price correction can *create* a spurious negative relationship between the two variables (for example, see Figure 3 results with Experimental Design 2, *σ*^2^ = 1 and *s*^2^ = 0.01).

If phenotypic plasticity evolves via the Unlinked Phenotypes mechanism (phenotypes in each environment evolve directly and independently of one another), neutral evolution will yield a negative relationship between baseline phenotype and reaction norm slope (Figure 4 “true” values). Furthermore, the negative relationship between these traits is maintained in the presence of measurement error (Figure 4 “estimated” values). In this case, however, adjusting the data using the Kelly and Price correction mostly eliminates the true negative relationships between the traits (Figure 4 “corrected” estimates). Conversely, if data are collected using Experimental Design 3, the true negative relationship between the traits is largely maintained. Nonetheless, as measurement error is increased the capacity for Experimental Design 3 to yield the true negative relationship decreases (Figure 4).

**Figure 4.**
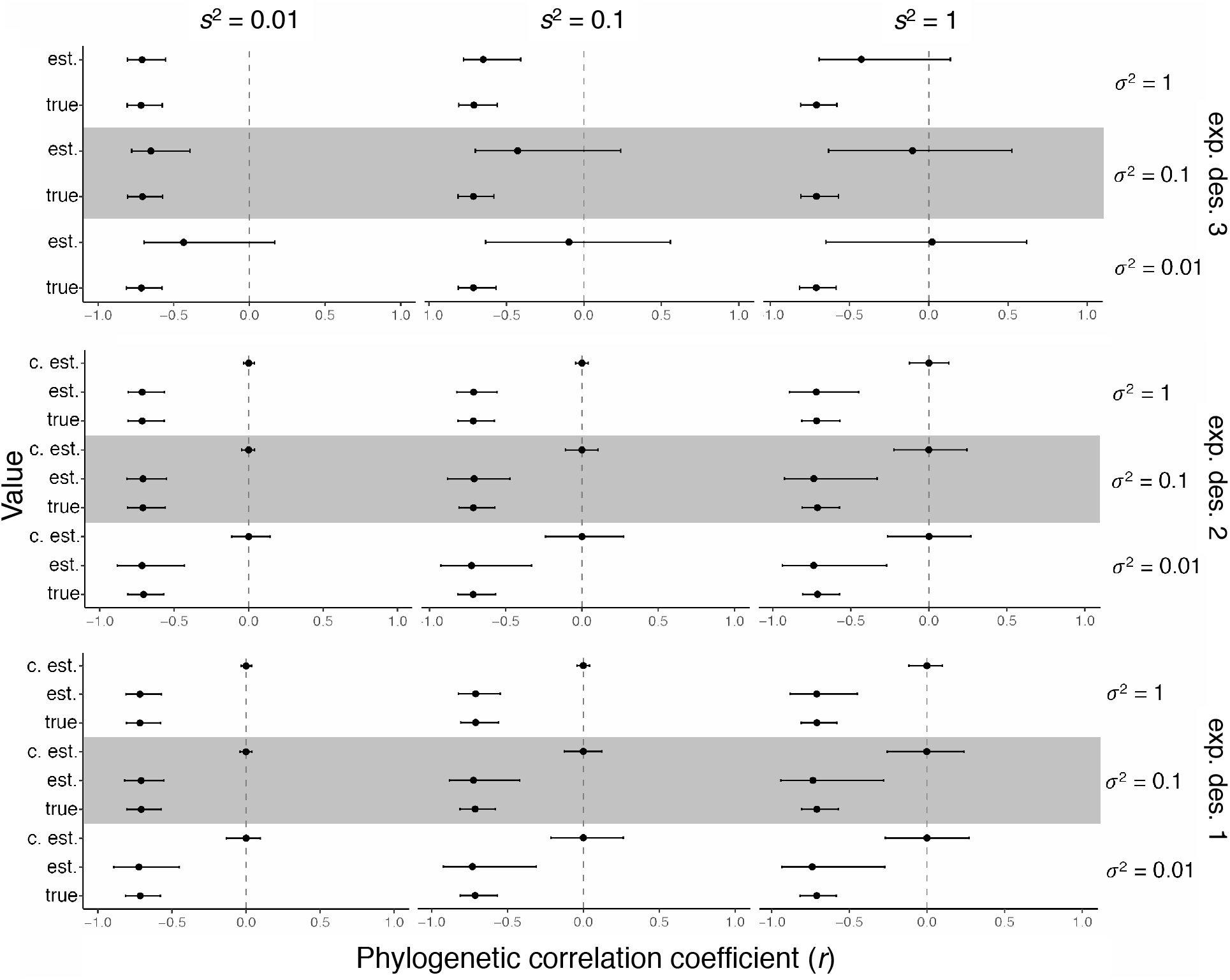
Summary of phylogenetic correlation coefficients between baseline phenotype and reaction norm slope from simulations in which plasticity evolved by Brownian motion via the Unlinked Phenotypes mechanism. Each panel shows the result of 1000 simulations. Each row shows the results of simulations with a different experimental design (Figure 1): *Bottom*, Design *Middle*: Design 2. *Top*: Design 3. Alternating shaded areas differentiate simulations with different Brownian motion rate parameters (*σ*^2^). Each column contains simulations with a different error variance for phenotypic estimation (*s*^2^). “true” denotes the true phylogenetic correlation values from the simulations. “est.” denotes estimates of the true correlation coefficients given measurement error. “c. est” denotes estimated correlation coefficients adjusted to remove the effect of regression to the mean following our modified Kelly and Price (2005) correction. Adjusted results are not given for Experimental Design 3 because the method cannot be applied to data collected with that design.

### Reanalysis of Nudibranch heat tolerance data

We re-analyzed data on heat tolerance plasticity in Nudibranchs (Armstrong et al. 2019) by applying our phylogenetic adaptation of the Kelly and Price (2005) correction. In the original analysis, the authors found a significant negative relationship between independent contrasts for baseline heat tolerance and heat tolerance plasticity (P < 0.005, Armstrong et al. 2019). In our re-analysis, we found that a negative relationship remained, but was no longer significant at the *σ* = 0.05 level (P = 0.069; Figure 5). In addition, the variance explained dropped from 88% to 51%. The maintenance of a negative relationship and relatively high explanatory power suggests that there may be a real biological relationship between baseline tolerance and plasticity in this group, but the need to adjust for regression to the mean raises the bar for the evidence required.

**Figure 5.**
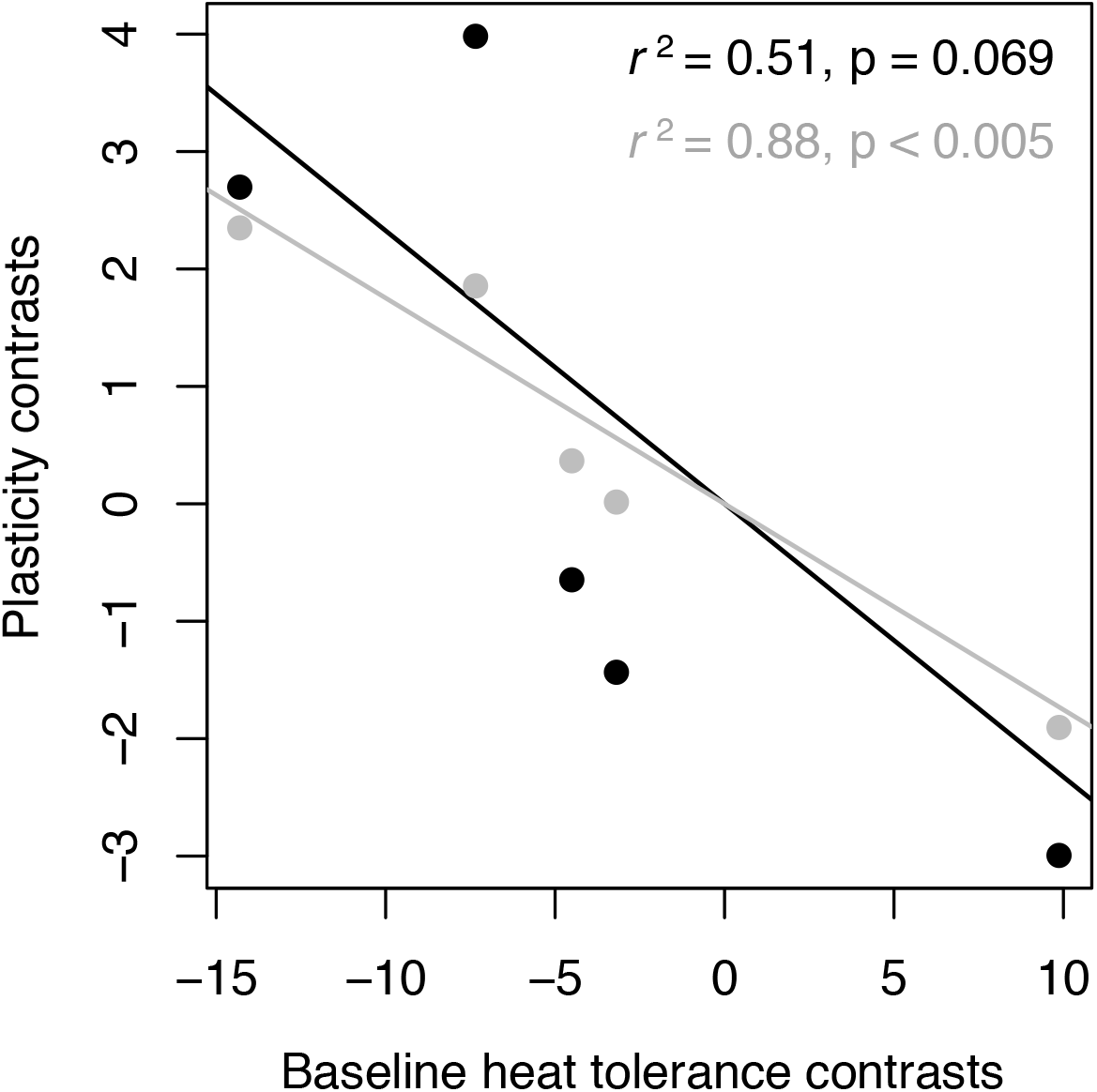
Nudibranch heat tolerance plasticity data from Armstrong et al. (2019). Phenotypes were collected using Experimental Design 2 (Figure 1). Original data and analysis in gray. Re-analysis applying the modified Kelly and Price (2005) method to remove regression to the mean effects in black.

## Discussion

Herein, we have identified three general patterns important for testing and understanding the evolution of genetic assimilation at the macroevolutionary level: 1) regression to the mean is a pervasive problem; 2) experimental design and data correction can minimize the effect of regression to the mean; and 3) neutral evolutionary processes can produce patterns consistent with genetic assimilation, depending on the linkage between trait expression in different environments. Below, we discuss each of these findings in greater detail.

### Regression to the mean when plasticity does not diverge

When phenotypic plasticity was present but did not diverge among taxa, measurement error alone generated spurious correlations between baseline phenotype and plasticity under Experimental Designs 1 and 2 (Figure 2 “estimated” values). Furthermore, the magnitude of the correlation coefficient was affected very little by the magnitude of measurement error simulated (Figure 2). This is because error variation cannot be swamped by biological variation when there are no phenotypic differences among taxa. These results clearly demonstrate that regression to the mean can be a significant problem when testing for genetic assimilation at the macroevolutionary level. The choice of experimental design, however, also matters considerably. In particular, when data are collected using Experimental Design 3, regression to the mean is not a persistent problem (Figure 2).

Why are Experimental Designs 1 and 2 susceptible to regression to the mean while Experimental Design 3 is not? The crux of the matter is that with Designs 1 and 2, the reaction norm slope depends on the estimated baseline phenotype. Designs 1 and 2 have only one estimate of baseline phenotype. That single baseline phenotype is also used to estimate plasticity. If measurement error causes an estimate of baseline phenotype to be above the true value, that will drive down the estimated reaction norm slope. Conversely, if error causes a baseline estimate to be below its true value, this will in turn drive up the estimated reaction norm slope. With Design 3, this cannot happen because the baseline phenotype and reaction norm are estimated using separate samples.

Design 3 demands two independent estimates of phenotype under baseline conditions, which we will call e1 and e2. One of those estimates (let’s say e1) is used only as the baseline phenotype, but does not contribute to the calculation of plasticity. The other estimate (e2) contributes only to the calculation of plasticity. Under this design, if measurement error causes e1 to be above or below the true value it will have no effect on the plasticity estimate, which depends only on e2.

All hope is not lost with Experimental Designs 1 and 2, however. Using our modified Kelly and Price correction, phenotypic values can be adjusted such that the bias towards a spurious negative relationship is removed (Figure 2 “corrected” values; though see below for caveats). Next, we discuss the implications of these different approaches when plasticity itself evolves.

### Regression to the mean under neutral evolution of plasticity (Linked Phenotype model)

Under the Linked Phenotypes model, the intercept (essentially, the baseline phenotype) and reaction norm slope evolve via Brownian motion. Under this model, neutral evolution does not produce a negative relationship between baseline phenotype and plasticity under any experimental design (Figure 3 “true” values). The introduction of measurement error, however, can again lead to regression to the mean under Experimental Designs 1 and 2 (Figure 3 “estimated” values). The degree to which regression to the mean is a problem depends on the magnitude of measurement error relative to the variance in true trait values among taxa as denoted by the Brownian motion rate parameter. If measurement error is large and/or phenotypic variation among taxa is low, regression to the mean emerges consistently for Experimental Designs 1 and 2. By contrast, if measurement error is small and phenotypic variation among taxa is large, regression to the mean is much less of an issue (Figure 3 “estimated” values).

Applying the Kelly and Price correction to data collected under Experimental Designs 1 and 2 can remove the effect of regression to the mean when it is present (Figure 3 “corrected” values). Our results, however, also highlights the caveat that a Kelly and Price correction should only be applied if the initial analysis yields a significant negative relationship. When initial analyses yield no significant negative relationship, applying the Kelly and Price correction can itself create a spurious negative relationship between baseline phenotype and plasticity. Experimental Design 3 again avoids the problem of regression to the mean due to the estimation of two baseline phenotypes (see above).

### Regression to the mean under neutral evolution of plasticity (Unlinked Phenotypes model)

Under the Unlinked Phenotypes model, the phenotypes in each environment evolve independently of one another, and the magnitude of plasticity emerges as a byproduct of the difference between the two phenotypic values. Under these conditions, a true negative association between baseline phenotype and plasticity consistently arises under Brownian motion evolution (Figure 4 “true” values). In other words, there are conditions in which neutral evolution alone can produce a pattern consistent with genetic assimilation. This is in contrast to the most common explanations for genetic assimilation, which invoke some combination of selection and/or genetic constraint to explain why a reduction in plasticity evolves (West-Eberhard 2003; Pigliucci et al. 2006; Ehrenreich and Pfennig 2016).

Unlike the Linked Phenotypes model, experimental error has little to effect on the correlation coefficients calculated with the Unlinked Phenotypes model using Experimental Designs 1 and 2 (Figure 4 “estimate” values). The reason for this is that phenotypic evolution via Brownian motion in the Unlinked Phenotypes model amounts to random independent increases or decreases in the phenotypes expressed in each environment, while experimental error leads to spurious random increases or decreases in the phenotypes measured in each environment, which from a statistical standpoint is the same process.

When data from the Unlinked Phenotypes model are adjusted using the Kelly and Price correction, the negative relationship between baseline phenotype and plasticity disappears (Figure 4, “corrected” values). In this case, however, the correction does not remove a spurious relationship due to regression to the mean. Instead, it removes a true relationship that resulted from a neutral evolutionary process. This occurs because of the conceptual foundation upon which the Kelly and Price correction is predicated: it assumes that a hallmark of regression to the mean is that phenotypic values between treatments (in this case, environments) within experimental units are uncorrelated with one another. This, of course, is the precise pattern that neutral evolution will produce if the phenotypes induced by plasticity are unlinked. Our finding highlights that biological variation and error variation can be difficult to separate (Kelly and Price 2004; Hansen and Bartoszek 2012), and statistical correction for regression to the mean is not a panacea.

Correlation coefficients estimated using Experimental Design 3 can maintain the true negative relationship between baseline phenotype and plasticity, but only if the measurement error is not too large. As measurement error grows relative to the true phenotypic variance among taxa, estimated correlation coefficients move away from the true values and towards 0 (Figure 4).

### Implications

Our results make clear that regression to the mean is a serious statistical problem that can confound tests of genetic assimilation at macroevolutionary scales. Whether or not regression to the mean is an issue in any particular study will depend on the experimental design used. Unfortunately, the problematic experimental designs (our Designs 1 and 2) are those most often used to test for plasticity and genetic assimilation. This is certainly true within the field of thermal physiology, where tests of the Trade-Off Hypothesis, which states that greater thermal tolerance evolves via genetic assimilation (Heerwaarden and Kellerman 2020), are almost invariably based on data collected using Experimental Designs 1 and 2 (e.g., Armstrong et al. 2019). These studies typically find support for the Trade-off Hypothesis but do not account for regression to the mean.

Some evolutionary physiologists have used an alternative approach to test the Trade-off Hypothesis, in which thermal tolerance plasticity is correlated with habitat temperature (van Heerwaarden and Kellermann 2020). This approach avoids the regression to the mean problem, but it is important to recognize that modeling the association between plasticity and habitat tests a different hypothesis than modeling the association between plasticity and baseline phenotype. The latter directly pertains to the evolution of within-individual trait associations, trade-offs, and genetic assimilation. The former does not.

Fortunately, data collected using Experimental Designs 1 and 2 can be successfully adjusted to remove regression to the mean using the Kelly and Price method. For example, when plasticity is invariant among taxa, adjusting the data removes spurious relationships between baseline phenotype and plasticity due to measurement error (Figure 3). The same holds under neutral evolution when there is linkage between the traits expressed in each environment (Figure 3). The caveat is that adjusting data using the Kelly and Price method can incorrectly obfuscate a true negative relationship between baseline phenotype and plasticity if the traits expressed in each environment are unlinked (Figure 4).

The need to adjust data can be avoided by using Experimental Design 3. This approach does not lead to spurious negative relationships between baseline phenotype and plasticity (Figures 3 and 4). It also recovers true negative relationships when they occur (Figure 4). Unfortunately, however, this experimental design is significantly more labor intensive that Designs 1 and 2. It requires that we estimate phenotypes on more groups of individuals (3 versus 2), and its reliability depends on measurement error in the estimation of phenotypic trait means for each environment, making large sample sizes crucial. There are issues with Experimental Design 3 as well. For example, when individuals have their phenotypes measured more than once, carryover effects of previous experience can influence results (O’Connor et al. 2014). Furthermore, for some taxa and traits, measuring the same individual twice will simply not be possible. One solution in such cases is to have three groups of individuals per taxa: two that have their phenotype measured under baseline conditions, and one that has phenotype measured under novel conditions. One baseline group is used to calculate plasticity, the other is used as the baseline phenotype estimate: the key is that different baseline measurements are used for each.

Whether or not induced phenotypes are genetically linked influences null expectations for reaction norm evolution. Specifically, a macroevolutionary pattern consistent with genetic assimilation can arise under a neutral evolutionary process in the absence of selection and genetic linkage between phenotypes. The importance of genetic linkage for reaction norm evolution has long been appreciated. For example, the classic work of Via and Lande (1985) demonstrated that genetic covariance between induced phenotypes influences the trajectory of reaction norm evolution under selection. Unfortunately, genetic linkage between environmentally-induced phenotypes is almost certainly trait, population, and environment dependent, and in many instances knowledge about linkage is completely lacking. Our results highlight the need for greater scrutiny of the genetic architecture of induced phenotypes.

Overall, we show that great care is necessary both in the design of macroevolutionary tests for genetic assimilation, as well as in the interpretation of our results. A lack of correlation between plasticity and baseline phenotype is not the correct null hypothesis in many cases due to the statistical phenomenon of regression to the mean. Once this is recognized, the potential solutions to the regression to the mean problem must be applied with care. Additionally, the expected null hypothesis under neutral evolution depends on the mechanism that underlies the expression of phenotypic plasticity. We hope that this analysis will be a helpful reference for others in the design and analysis of future studies of hypothesized genetic assimilation in phenotypically plastic traits.

## Data availability

Data and code to reproduce all the analyses of this study available at: GitHub: https://github.com/liamrevell/Gunderson-and-Revell-2021.

## Notes

### Competing Interest Statement

The authors have declared no competing interest.

