## Supplementary Figure S1 for "Testing for genetic assimilation with phylogenetic comparative analysis: Conceptual, methodological, and statistical considerations"

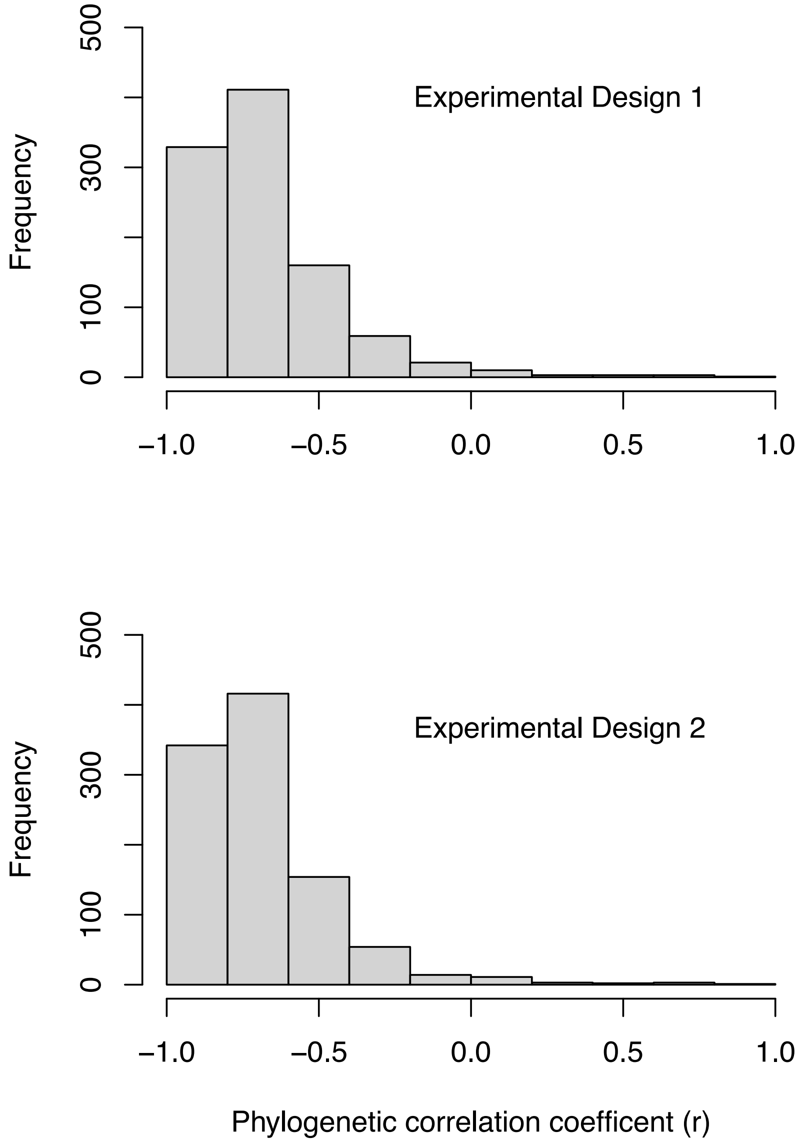


Supplementary Figure S1. Distribution of phylogenetic correlation coefficients from simulations in which there was no plasticity, plasticity does not evolve, and phenotypes in each environment are measured with error with s.d. = 0.1. N = 1000 simulations per plot. Top: simulations with Experimental Design 1. Bottom: simulations with Experimental Design 2. See main text for general details about how simulations were constructed and the different Experimental Designs.
